# A simple strategy to reduce the salivary gland and kidney uptake of PSMA targeting small molecule radiopharmaceuticals

**DOI:** 10.1101/2020.07.24.220277

**Authors:** Teja Muralidhar Kalidindi, Sang-Gyu Lee, Katerina Jou, Goutam Chakraborty, Myrto Skafida, Scott T. Tagawa, Neil H. Bander, Heiko Schoder, Lisa Bodei, Neeta Pandit-Taskar, Jason S. Lewis, Steven M. Larson, Joseph R. Osborne, Naga Vara Kishore Pillarsetty

## Abstract

The past five years have seen an increasing acceptance of peptide-based prostate-specific membrane antigen (PSMA)-targeted radionuclide therapy (TRT) agents for treatment of metastatic castration-resistant prostate cancer (mCRPC), with [^177^Lu]-DKFZ-PSMA-617 ([^177^Lu]-PSMA-617) emerging as the leading candidate. [^177^Lu]-PSMA-617 and other PSMA ligands have shown efficacy in reducing the tumor burden in mCRPC patients but irradiation to salivary gland and kidneys is a concern and dose limiting factor. Therefore, methods to reduce non-target organ toxicity are needed to safely treat patients and preserve their quality of life. Here, we report the effects of the addition of the cold PSMA ligand DKFZ-PSMA-11 (PSMA-11) on the uptake of [^177^Lu]-PSMA-617 in tumor, salivary glands and kidneys. Groups of athymic nude mice (n = 4) bearing PC3-PIP (PSMA+) tumor xenografts were administered with [^177^Lu]-PSMA-617 along with 0, 5, 100, 500, 1000 and 2000 pmoles of PSMA-11. Biodistribution studies 1 h post-administration revealed that [^177^Lu]-PSMA-617 uptake in PSMA-expressing PC3-PIP tumors was 21.71±6.13, 18.7±2.03, 26.44±2.94, 16.21±3.5, 13.52±3.68, and 12.03±1.96 %ID/g when 0, 5, 100, 500, 1000 and 2000 pmoles of PSMA-11 were added, respectively. Corresponding kidney uptake values were 123.14±52.52, 132.31±47.4, 84.29±78.25, 2.12±1.88, 1.16±0.36, 0.64±0.23 %ID/g, respectively. Corresponding salivary gland uptake values were 0.48±0.11, 0.45±0.15, 0.38±0.3, 0.08±0.03, 0.09±0.07, 0.05±0.02 % ID/g, respectively. Thus, uptake of PSMA TRT agents in salivary gland and kidney can be substantially reduced without impact on tumor uptake by adding cold PSMA-11. Our data provides proof-of-concept and we propose that similar strategy be pursued in future clinical trials to prevent xerostomia and renal toxicity arising from [^177^Lu]-PSMA-617.

## Introduction

In 2020, an estimated 192,000 new patients will be diagnosed with prostate cancer in US ^1^, adding to the 3.6 million men who have been previously diagnosed with prostate cancer (PC). Of those new patients, 33,330 (∼17%) will die from metastatic disease, despite the implementation of therapies such as androgen deprivation therapy (ADT), surgery, immunotherapy, chemotherapy and radiation therapy ^1^. To overcome the limitations of current treatment strategies, researchers are developing novel therapies, including targeted radionuclide therapy (TRT) with therapeutic radionuclides (iodine-131, lutetium-177, actinium-225, thorium-227) appended to molecules that target PC cells or their microenvironment ^2^. Prostate specific membrane antigen (PSMA), also known as folate hydrolase-1 (FOLH1), is an androgen receptor (AR)-regulated target gene that is highly overexpressed on the apical membrane of both localized and metastatic PC lesions, especially in non-responding patients ^3^. Transcriptomic analysis of tumor biopsy samples of PC patients clearly indicates that the expression of PSMA is upregulated at transcript level with the progression of disease (**Fig 1**) ^4^. Because PSMA is highly expressed in PC ^5-7^, provides an attractive target, and can even be used to treat lesions that stop responding to conventional therapies, it has become a subject of intense interest to the nuclear medicine clinical community, both as an imaging and a therapeutic agent.

**Figure 1:**
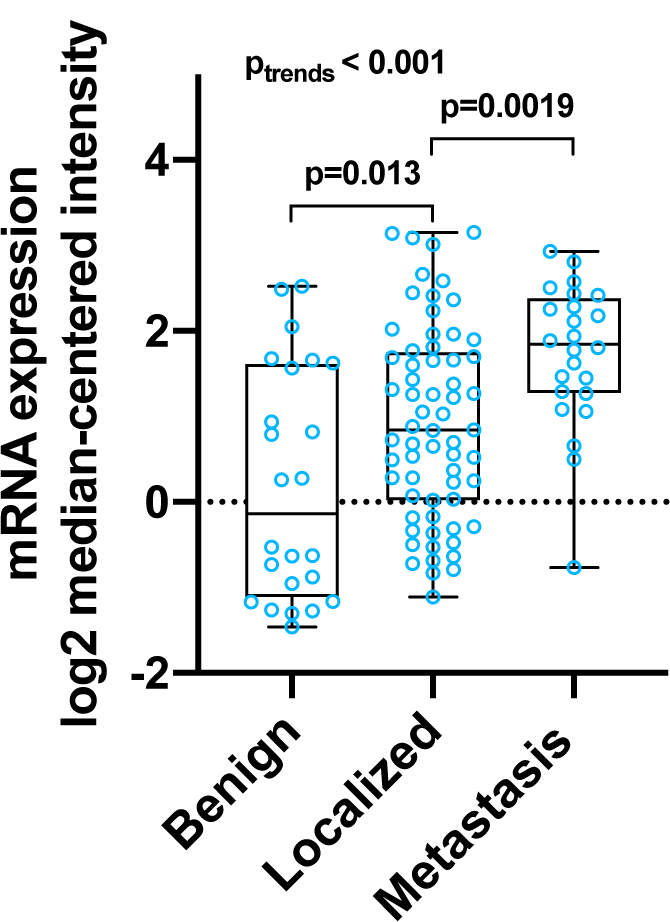
PSMA mRNA levels in prostate cancer samples analyzed from the Yu prostate data set (4). A total of 112 samples including 23 normal prostate, 64 prostate carcinoma, and 25 metastatic prostate cancer samples were analyzed on Affymetrix U95A-C microarrays; 8603 gene transcripts were analyzed and the PSMA (FOLH1) mRNA expression data was plotted using Graphpad Prism. P value calculated by unpaired student t test. P-trends were analyzed by one-way ANOVA.

The potential of PSMA as a target for imaging of prostate cancer was initially demonstrated with imaging agent [^111^In]-Capromab Pendetide (Prostascint®) in 1996 ^8^. Independently, Bander’s group developed the humanized monoclonal antibody J591, which targets the external epitope of PSMA. Imaging studies with radiolabeled J591 demonstrated its superiority over [^111^In]-Prostascint® using PET and SPECT imaging techniques ^9, 10^. J591 radiolabeled with therapeutic isotopes (^90^Y, ^177^Lu and ^225^Ac) has demonstrated efficacy both in preclinical and clinical studies ^11, 12^. However, due to the long plasma circulation time of the antibodies, a dose-limiting myelotoxicity was observed.

To circumvent the problems arising from the long plasma half-life of circulating mAbs, alternative PSMA-targeting agents with significantly shorter plasma half-life have been developed. These include small molecules ^13^, affibodies, minibodies ^14^, single chain fragments ^15, 16^. Most of the small-molecule-based PSMA-targeting radiopharmaceuticals have a common glutamyl-urea-amino acid moiety (indicated in red in **Fig. 2**) as initially proposed by Pomper et. al. in 2005 ^17^. These radiopharmaceuticals bind to the catalytic site of PSMA with low nanomolar affinities; because of their low molecular weight (< 2000 Da) and hydrophilic nature, they rapidly clear from the blood pool. This results in substantially lower radiation doses to bone marrow/hematopoietic system can be achieved ^18^. This has led to the rapid clinical translation of many glutamyl-urea-amino acid derivatives both for diagnostic and therapeutic applications in prostate cancer ^19^. As of April 2020, a simple search in the clinicaltrials.gov website reveals more than 225 clinical trials on radiolabeled small molecules targeting PSMA in prostate and other cancers. While several different agents are in clinical trials, the most widely studied include [^68^Ga]-DKFZ-PSMA-11, [^18^F]-DCPyFl, [^131^I]-MIP-1095, [^177^Lu]-DKFZ-PSMA-617, [^225^Ac]-DKFZ-PSMA-617, [^177^Lu]-PSMA-I&T ^20^. The efficacy of these small molecule-based imaging agents, especially [^68^Ga]-DKFZ-PSMA-11, in identifying metastatic disease foci has been established ^21^. Imaging studies with [^68^Ga]-DKFZ-PSMA-11 revealed high uptake in tumors and moderate-to-high uptake in normal organs that express PSMA, including salivary and lacrimal glands, small bowel, and kidneys ^3^. On the therapeutic front, the first-in-human studies were performed using the radioiodinated derivatives [^124/131^I]-MIP-1095 from the Heidelberg group ^22^. Targeting was assessed using PSMA PET/CT and SPECT imaging studies that indicated excellent tumor uptake, moderate uptake in liver and proximal intestine, and high uptake in salivary glands, lacrimal glands and kidneys is related to high expression of PSMA in these organs ^23^. The mRNA and protein expression data from the human protein atlas website also confirms the presence of PSMA at varying levels in normal prostate, small bowel (duodenum), kidney and salivary glands ^24^. Dosimetry estimates for [^124/131^I]-MIP-1095 revealed that the tumors received doses up to 6.25 mSv/MBq. Apart from the tumor, the highest absorbed doses were delivered to the salivary glands (3.8 mSv/MBq), with minimal radiation dose to the red marrow (0.37 mSv/MBq). More than 60% of patients demonstrated prostate-specific antigen (PSA) decline. Owing to high tumor uptake and background clearance [^177^Lu]-DKFZ-PSMA-617 is currently in phase 3 clinical trial. Dosimetry studies revealed that tumors received on average 3.3 mGy/MBq, leading to > 50% decline in PSA in 55% of selected patients ^25^. These findings led to the rapid expansion of the use of [^177^Lu]-PSMA-617 in clinical trials as a salvage option for patients who had stopped responding to traditional hormonal therapies, [^223^Ra]-RaCl_2_, chemotherapy and immunotherapy. Common toxic side effects included grade 1–2 dry mouth, transient nausea, and fatigue; grade 3-4 thrombocytopenia was observed rarely (< 15%). While no prospective study has been completed, retrospective reports of [^225^Ac]-PSMA-617 demonstrate significant responses even in patients who are refractory to [^177^Lu]-PSMA-617 therapy ^26^. Although, myelotoxicity was mild, grade 1–2 xerostomia was commonly observed (87% of patients) ^25^. This can be expected because salivary glands are one of the dose-limiting toxicity organs receiving a dose of about 1 Gy/GBq of [^177^Lu]-PSMA-617 ^27^. The other dose-limiting organ are kidneys, which receive a dose of about 0.5–0.6 Gy/GBq of [^177^Lu]-PSMA-617 and in which mild grade 1–2 renal dysfunction is observed. The salivary gland toxicity and possible renal toxicity can limit the scope of usage of PSMA agents particularly when labelled with alpha-emitters, such as [^225^Ac]-PSMA-617, despite their tremendous potential to improve and extend the lives of mCRPC patients. Therefore, there is a clear and urgent unmet need to develop methods that can minimize exposure to the salivary gland and kidney and ensure the success of PSMA targeted radiotherapies.

**Figure 2:**
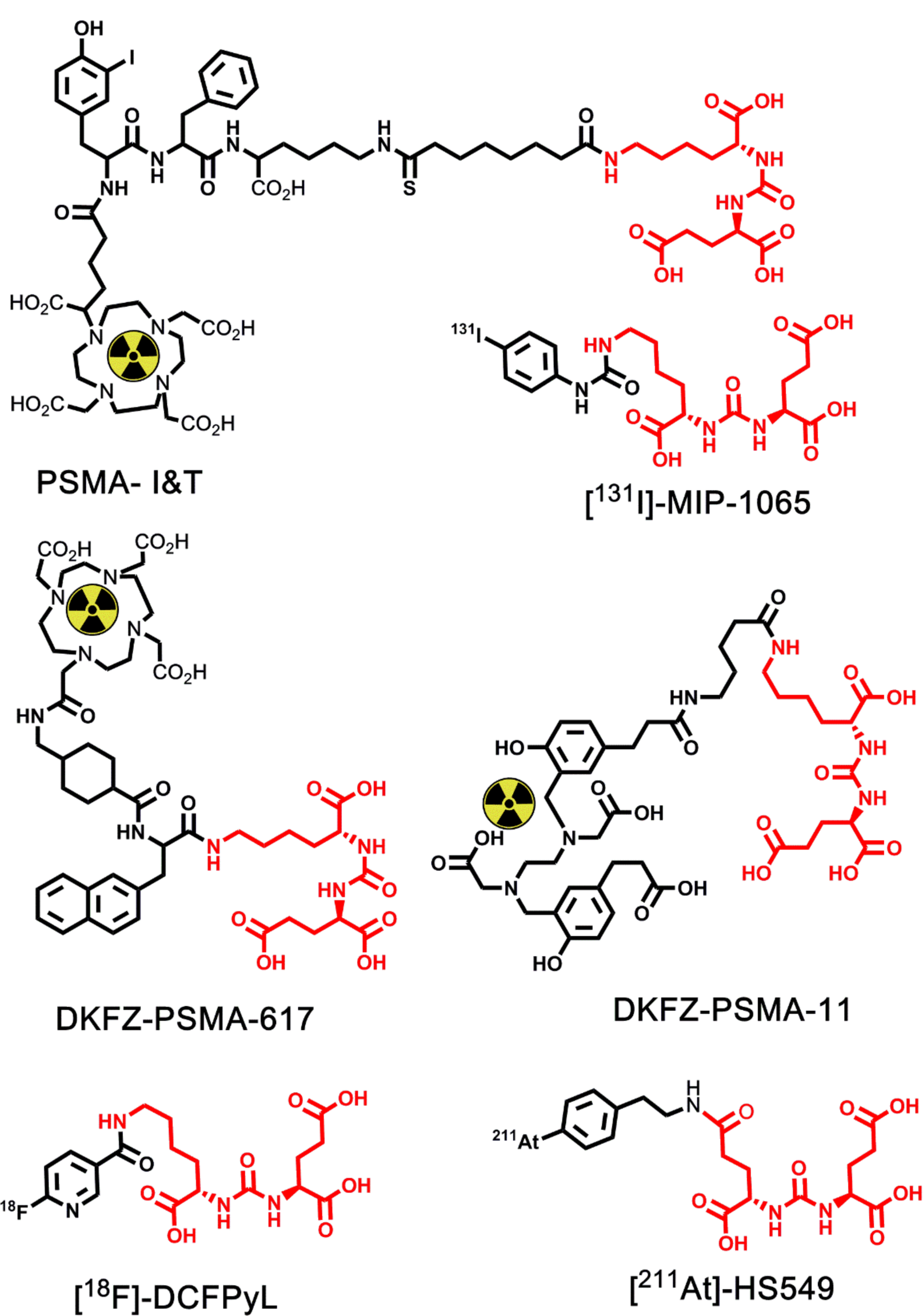
Representative structures of peptide-based PSMA-targeted radiopharmaceuticals currently being evaluated in several preclinical and clinical studies.

In our previously reported work with [^68^Ga]-PSMA-11 in mice models, we observed that adding excess of the ligand DKFZ-PSMA-11 to the radiopharmaceutical [^68^Ga]-PSMA-11 reduces the uptake of [^68^Ga]-PSMA-11 in salivary glands and kidneys without significantly affecting the uptake in tumor ^28, 29^. Addition of the ligand results in the reduction of the effective molar activity (EMA), which is defined as the activity of the targeted radiopharmaceutical divided by the total amount of the targeted agent (radiolabeled pharmaceutical + free ligand) in moles. This finding led us to hypothesize that reducing the effective molar activity of the therapeutic radiopharmaceutical [^177^Lu]-PSMA-617 could be used to reduce the accumulation of radiopharmaceutical in the salivary glands and kidneys, thereby decreasing non-target organ toxicity. In the current report, we present the results of our study on the uptake of [^177^Lu]-PSMA-617 in the tumor, salivary gland and kidney of athymic nude mice bearing PC3-PIP xenografts as a function of reducing the effective molar activity of our radiopharmaceutical by addition of cold ligand PSMA-11.

## Methods

All starting materials, solvents, and reagents were purchased from commercial sources (Sigma Aldrich, Fisher Scientific, ABX or MedKoo Biosciences Inc.) and used without further purification. The radiochemical precursor DFKZ-PSMA-617 (catalog number 206934) was purchased from MedKoo Biosciences Inc. (Morrisville, NC) and dissolved in metal free water to achieve a concentration of 1 mg/mL and 5 µg (5 µL) was aliquoted in 1.5 mL microcentrifuge tubes and stored at −80 °C and used for complexation reactions. DKFZ-PSMA-11 (catalog number 9920) was obtained from ABX advanced biochemical compounds GmbH (Radeberg, Germany). DKFZ-PSMA-11 (PSMA-11) was dissolved in metal-free water to achieve a concentration of 1 mg/mL and used for reducing the effective molar activity. All solvents used for HPLC analysis and purification within this project were purchased from Fisher Scientific (HPLC grade). For radiosynthesis TraceSELECT™ grade sodium acetate was used. Water (>18.2 MΩ cm-1 at 25 °C) was obtained from an Alpha-Q Ultrapure water system from Millipore (Bedford, MA) and used for purification and HPLC purposes. Optima® grade acetonitrile was purchased from Fisher Scientific (Hampton, NH) and was used for HPLC purposes. Trifluoroacetic acid was purchased from Sigma Aldrich. Purification cartridge Strata™-X (#8B-S100-TAK, 33µm Polymeric Reversed Phase C-18, 30 mg/1mL) and analytical HPLC column Luna (00G-4252-E0, 250 x 4.6 mm, 5 µ, 100 A°, C-18 (2) with TMS end capping) were purchased from Phenomenex (Torrance, CA). Lutetium-177 in the form of lutetium chloride (^177^Lu-LuCl_3_) was obtained from Missouri University Research Reactor (MURR, Columbia, MO). Radioactivity was measured using Wizard 2480 gamma counter (Perkin Elmer, Waltham, MA).

### Cell Lines

PC3-PIP cells (PC3-PSMA-IRES-Puromycin; kindly provided by Dr. Martin Pomper – Johns Hopkins University, Balitomore MD) were maintained under RPMI+10%FCS (Fetal Calf Serum)+PS (Penicillin and Streptomycin) media, puromycin, penicillin-streptomycin and used for our current studies.

### Prostate Cancer Xenograft model

PC3-PIP cells were grown in RPMI+10%FCS (Fetal Calf Serum)+PS (Penicillin and Streptomycin) media. 5 million cells were used per xenograft. The cells were prepared in 200 μl for xenografts in a solution of media and Matrigel® in a 1:1 ratio and were injected subcutaneously. The mice were monitored weekly, and by Week 3 the tumors reached an average size of 200–300mm^3^, at which point they were used for the experiment.

### Radiosynthesis

Lutetium-177 in the form of lutetium chloride (^177^Lu-LuCl_3_) was obtained from Missouri University Research Reactor (MURR, Columbia, MO). 30 µL of 0.25 M sodium acetate buffer (pH 5.5) was added to a 1.5 mL Eppendorf tube containing 5 nmoles of DKFZ-PSMA-617 (5 µl, 1 mM solution in water) and briefly vortexed. To the resulting solution, 0.7 µL of [^177^Lu]-LuCl_3_ (1.5 mCi) was added, then centrifuged for 20 seconds (500 rpm) and heated to 95 °C for 30 min to yield crude [^177^Lu]-DKFZ-PSMA-617. The reaction mixture was allowed to cool and loaded onto a C18 reverse-phase cartridge (Strata(tm)-X cartridge; Cat No. 8B-S100-TAK, 33 µm Polymeric Reversed Phase C-18, 30 mg/1 mL) that was preconditioned with 1 mL of 95% ethanol followed by 2.5 mL of pure water. The cartridge was then rinsed with 0.5 mL of water to remove non-chelated [^177^Lu]-LuCl_3_. The pure product (1.45 mCi) was eluted using 450 µL of 66% ethanol in 0.9% saline solution. The molar activity of the product was 0.29 mCi/nmole.

### Animal Doses

For formulating doses with reduced effective molar activity, calculated amounts of PSMA-11 (1 mg/mL in water) were added to the formulation. About 1.45 μCi of [^177^Lu]-DKFZ-PSMA-617 at specific activity of 0.29 mCi/nmole containing about 5 pmoles of PSMA-617 was used directly or diluted with PSMA-11. Individual doses of [^177^Lu]-DKFZ-PSMA-617 containing 0, 5, 100, 500, 1000 and 2000 pmoles of PSMA-11 were prepared to yield [^177^Lu]-DKFZ-PSMA-617 with effective molar activities of 0.29, 0.145, 0.0138, 0.00287 0.00144 and 0.000723 mCi/nmole. Considering average mass of mice (25 g) this translates to 0, 0.2, 4, 20, 40 and 80 nmoles of PSMA-11/kg dose in addition to the standard [^177^Lu]-DKFZ-PSMA-617 dose.

### Biodistribution Studies

[^177^Lu]-DKFZ-PSMA-617 (7.5 µCi, 2.75 MBq) containing 0, 5, 100, 500, 1000 and 2000 pmoles of PSMA-11 was administered via tail vein to different cohorts (n = 4) of athymic nude mice bearing PC3-PIP xenografts. Activity and weights of the syringes were measured pre- and post-injection to calculated amount of administered radioactivity. The mice were sacrificed by CO_2_ asphyxiation 1 h post-administration. Blood was collected immediately post-sacrifice by cardiac puncture and collected in a pre-weighed tubes. Necropsy was performed to collect tumor (PC3-PIP), heart, lungs, liver, spleen, stomach, small intestine, large intestine, kidney, muscle bone, tail and salivary glands. The organs were washed with water, air dried, transferred into pre-weighed tubes and counted for radioactivity on gamma counter. For calculating administered dose, several standards were prepared and counted along with the organs. Total administered counts were determined based on the difference in weight before and after injection of the activity. The counts from the gamma-counter were divided by injected counts. For each sample obtained, count data was background- and decay-corrected and the tissue uptake was measured in units of percent injected dose per gram (%ID/g) by dividing tissue counts to total administered counts normalized to the weight of the tissue multiplied by 100.

## RESULTS

### Synthesis of [^177^Lu]-DKFZ-PSMA-617

[^177^Lu]-DKFZ-PSMA-617 was synthesized in high purity (> 97%) with minimum molar activity of 11.1 GBq/µmole (300 mCi/µmole). No attempts to improve molar activity were made as this was enough for the intended application.

### Biodistribution Studies

Table 1 provides the results of our 1 h biodistribution study in mice bearing PC3-PIP tumors of radiopharmaceutical [^177^Lu]-DKFZ-PSMA-617 (55.5 kBq, 1.5 μCi, 5 pmoles) with addition of 0, 5, 100, 500, 1000 and 2000 pmoles of DKFZ-PSMA-11. The total amount of the PSMA-targeting ligand is 5, 10, 105, 505, 1005 and 2005 pmoles, resulting in formulation of effective molar activities of 0.29, 0.145, 0.0138, 0.00287, 0.00144 and 0.000723 mCi/nmol. As expected, when no PSMA-11 was added to the [^177^Lu]-PSMA-617 we observed high uptake in PSMA-expressing PC3-PIP tumor and kidneys and small but significant uptake in salivary glands. As the effective molar activity of the formulation was progressively decreased by addition of PSMA-11, we observed significant reductions in uptake of the radiopharmaceutical in the kidneys and salivary glands. Uptake of [^177^Lu]-PSMA-617 in PC3-PIP tumors was 21.71 ± 6.13, 18.7 ± 2.03, 26.44 ± 2.9, 16.21 ± 3.5, 13.52 ± 3.68, and 12.03 ± 1.96%ID/g when 0, 5, 100, 500, 1000 and 2000 pmoles of PSMA-11 was administered along with [^177^Lu]-PSMA-617 — equivalent to effective molar activities of 0.29, 0.145, 0.0138, 0.00287, 0.00144 and 0.000723 mCi/nmol. Corresponding values in kidney were 123.14 ± 52.52, 132.31 ± 47.4, 84.29 ± 78.25, 2.12 ± 1.88, 1.16 ± 0.36, 0.65 ± 0.23 %ID/g, respectively. In the PSMA-expressing salivary gland the uptake values were 0.48±0.11, 0.45±0.15, 0.38±0.3, 0.08±0.03, 0.09±0.07, 0.05±0.02 %ID/g, respectively. Notably, adding 2000 pmoles of PSMA-11 to [^177^Lu]-PSMA-617 (equivalent to reducing the effective molar activity by 166-fold — 0.29 to 0.000723 mCi/nmole) — lowered mean uptake value (%ID/g) in PSMA+ tumor by about 44.5%, whereas in the kidneys and salivary glands the reduction of effective molar activity by 166-fold lowered mean uptake values by about 99.5% and 89.5%, respectively. These trends can be easily visualized in the graph that plots uptake (%ID/g) trends on a log10 scale as a function of effective molar activity (**Fig 3**). These experiments establish the feasibility of significantly reducing the uptake of [^177^Lu]-PSMA-617 in salivary glands and kidneys.

**Table 1:** Biodistribution data of [^177^Lu]-DKFZ-PSMA-617 (5 pmoles) as a function of total ligand mass in athymic nude mice bearing PC3-PIP xenografts at 1 h post-administration.

| **[$^{177}\text{Lu}$]-DKFZ-PSMA-617 uptake in organs as function of effective molar activity at 1 h** |  |  |  |  |  |  |
| --- | --- | --- | --- | --- | --- | --- |
|  | **0.29**<br>**mCi/nmole** | **0.145**<br>**mCi/nmole** | **0.0138**<br>**mCi/nmole** | **0.00287**<br>**mCi/nmole** | **0.00144**<br>**mCi/nmole** | **0.000723**<br>**mCi/nmole** |
| **Blood** | 0.36 ± 0.06 | 0.4 ± 0.08 | 0.5 ± 0.54 | 0.12 ± 0.06 | 0.11 ± 0.08 | 0.08 ± 0.08 |
| **Tumor** | 21.71 ± 6.13 | 18.7 ± 2.01 | 26.44 ± 2.94 | 16.21 ± 3.53 | 13.52 ± 3.68 | 12.03 ± 1.96 |
| **Heart** | 0.26 ± 0.14 | 0.32 ± 0.08 | 0.28 ± 0.34 | 0.04 ± 0.01 | 0.27 ± 0.28 | 0.03 ± 0.02 |
| **Lungs** | 0.78 ± 0.16 | 0.89 ± 0.34 | 0.66 ± 0.6 | 0.08 ± 0.03 | 0.2 ± 0.12 | 0.13 ± 0.13 |
| **Liver** | 0.19 ± 0.06 | 0.17 ± 0.05 | 0.25 ± 0.24 | 0.06 ± 0.01 | 0.09 ± 0.02 | 0.05 ± 0.02 |
| **Spleen** | 1.53 ± 0.88 | 1.72 ± 0.95 | 0.82 ± 0.77 | 0.09 ± 0.01 | 0.16 ± 0.14 | 0.05 ± 0.02 |
| **Stomach** | 0.11 ± 0.03 | 0.09 ± 0.04 | 0.45 ± 0.53 | 0.14 ± 0.17 | 0.03 ± 0.01 | 0.19 ± 0.31 |
| **S Intestine** | 0.19 ± 0.14 | 0.1 ± 0.02 | 0.6 ± 0.62 | 0.14 ± 0.07 | 0.08 ± 0.04 | 0.13 ± 0.15 |
| **L Intestine** | 0.07 ± 0.02 | 0.09 ± 0.05 | 0.25 ± 0.18 | 0.1 ± 0.07 | 0.05 ± 0.02 | 0.19 ± 0.25 |
| **Kidneys** | 123.1 ± 52.5 | 132.3 ± 47.4 | 84.3 ± 78.2 | 2.12 ± 1.88 | 1.16 ± 0.36 | 0.64 ± 0.23 |
| **Muscle** | 0.16 ± 0.08 | 0.13 ± 0.03 | 0.19 ± 0.13 | 0.05 ± 0.06 | 0.05 ± 0.02 | 0.02 ± 0.01 |
| **Bone** | 0.16 ± 0.12 | 0.33 ± 0.1 | 0.31 ± 0.15 | 0.11 ± 0.09 | 0.14 ± 0.16 | 0.18 ± 0.3 |
| **Tail** | 0.87 ± 0.32 | 0.74 ± 0.16 | 2.16 ± 2.57 | 0.44 ± 0.27 | 0.36 ± 0.24 | 0.33 ± 0.36 |
| **Salivary Glands** | 0.48 ± 0.11 | 0.45 ± 0.15 | 0.38 ± 0.3 | 0.08 ± 0.03 | 0.09 ± 0.07 | 0.05 ± 0.02 |

**Figure 3.**
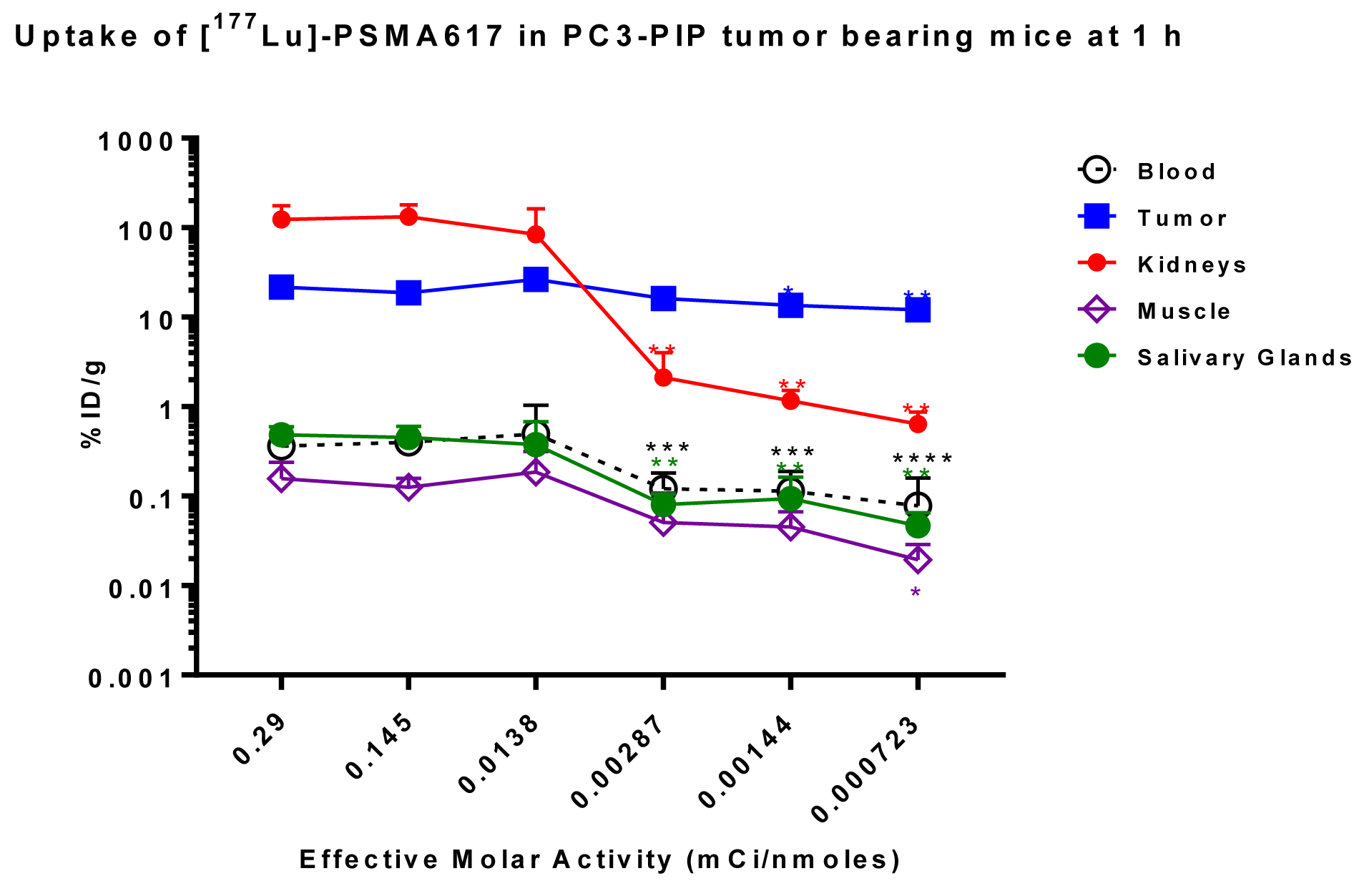
The absolute uptake of [^177^Lu]-PSMA-617 as a function of effective molar activity. The absolute uptake of [^177^Lu]-PSMA-617 in important organs at 1 h post-administration was plotted as a function of total mass of ligand administered to the animal. ANOVA was applied to analyze statistically whether effective molar activity affects absolute uptake of [^177^Lu]-PSMA-617. *P*-values for blood, tumor, kidney, and salivary glands were less than 0.001 and the p-value for muscle is 0.0145. After ANOVA, Dunnett’s multiple comparison test was performed using %ID/g at 5 pmoles of ligand as a control to analyze at what ligand mass the values differ statistically from the control group. *P*-value was marked in each group as asterisk (* *p* <0.05, ** *p* < 0.01, *** *p* < 0.005, **** *p* < 0.001). As shown in the figure, absolute uptake of [^177^Lu]-PSMA-617 in kidney, blood, and salivary gland is statistically lower when the total ligand mass is 500 pmoles or greater.

## Discussion

Small-molecule-based PSMA-targeted therapeutic radiopharmaceuticals have revolutionized the field of prostate cancer by providing treatment options to metastatic castration-resistant prostate cancer (mCRPC) patients who have exhausted all other treatment options. One agent ([^177^Lu]-PSMA-617) is the object of a currently ongoing randomized phase III registration trial. Both retrospective analysis of treatment data and prospective trials with therapeutic agents such as [^131^I]-MIP-1095, [^177^Lu]-PSMA-617, and [^225^Ac]-PSMA-617 have demonstrated significant therapeutic benefit in mCRPC patients ^19^. As intended, these small-molecule-based TRT agents targeting PSMA have proven able to reduce bone marrow toxicity ^22, 25^. Although cases of bone marrow toxicity grade 1 or higher were observed in a subset of patients undergoing PSMA TRT, it is likely that the patients experienced these side effects because either their bone marrow had been impoverished from prior therapies and/or they had high bone disease burden, which results in higher radiation dose to normal marrow next to the diseased sites.

PSMA expression in salivary glands and kidneys, however, poses significant challenges to the development and implementation of PSMA-targeted therapies, particularly for future applications with alpha-emitters. As a result of PSMA expression, patients have experienced grade 1–4 salivary gland toxicity, depending on the type of therapeutic agent. As expected these effects were particularly severe in patients being treated with [^225^Ac]-PSMA-617 ^26, 30^, while patients treated with [^177^Lu]-PSMA-617 experienced the relatively milder grade 1-2 toxicities ^25^. This represents a serious setback because the therapeutic responses with [^225^Ac]-PSMA-617 have been remarkablec even in mCRPC patients who failed to respond to [^177^Lu]-PSMA-617 ^26^. Though initial observations from several clinical studies indicated that renal toxicity has been mild to moderate with these agents ^27, 31^, due to the lack of specific binding or reuptake in the tubules, toxicity at later time points cannot be ruled out. Therefore, to ensure successful adoption of the small-molecule-based PSMA-targeted therapies, we must minimize non-target organ toxicity. Toward this goal, many groups have attempted to minimize radiation-induced damage to salivary glands, including monosodium glutamate co-administration ^32^, sialendoscopy with dilatation, saline irrigation and steroid injections ^33^, and external cooling of salivary and parotid glands ^34^. Regrettably, none of these techniques were particularly successful, while many proved to be rather cumbersome for routine clinical applications.

Our finding — that reducing the effective molar activity of PSMA-targeted small-molecule radiopharmaceuticals results in reduced salivary gland and kidney uptake — occurred during our work with the PSMA-targeted imaging agent [^68^Ga]-DKFZ-PSMA 11 ^28^. Encouraged by our results, we repeated these studies with the PSMA-targeting therapeutic radiopharmaceutical [^177^Lu]-DKFZ-PSMA-617 ^35^. Since multiple PSMA TRT agents are being developed, including [^131^I]-MIP-1095, [^177^Lu]-DKFZ-PSMA-617, [^177^Lu]-PSMA-I&T, [^225^Ac]-DKFZ-PSMA-617, [^177^Lu]-EB-PSMA-617, [^131^I]-MSK-PSMA1 etc., our goal was to develop a method that applies equally to all the therapeutics and diagnostics currently being developed ^7, 30, 35-38^. We reasoned that if we develop a method based on reducing molar activity by adding the specific cold ligand of the radiopharmaceutical (e.g. PSMA-617, EB-PSMA-617, PSMA-I&T), then the optimization must be done for each ligand separately because they have different pharmacokinetics. Therefore, the mass effect for each ligand is likely to be different in tumor and other organs. This represents a huge burden, as approval agencies might request clinical safety and efficacy data for each ligand separately. A major advantage of using PSMA-11 as a diluent to reduce effective molar activity is that its safety profile is well-established (at least in μg amounts) and, most importantly, it is not immunogenic. In addition, it is not protected by intellectual property, unlike most PSMA-targeting ligands. Therefore, if successful, it will permit immediate, international adoption of the technology across all groups working with the various PSMA TRTs — without the need for approval from patent/license holders. As such, we decided to use PSMA-11 as a diluent to reduce the effective molar activity of our preparation.

The mass of the preparation of [^177^Lu]-PSMA-617 administered to mice was in the range used for conducting clinical studies. Fendler et al. reported the results of [^177^Lu]-PSMA-617 study with the preparation’s molar activity being approximately 0.05 GBq/nmole (1.35 mCi/nmole) and Hoffman et al. used the [^177^Lu]-PSMA-617 preparation at a molar activity of 0.10 GBq/nmole (2.7 mCi/nmole) for their clinical trials ^25, 27^. Both groups used no carrier-added lutetium-177 for the preparation of their radiopharmaceuticals and the dose administered was about 6 GBq. Thus, assuming the average human weight to be about 70 kg, the total amount of the [^177^Lu]-PSMA-617 preparation administered was about 0.85 nmoles/kg (Hoffman et al.) to 1.7 nmoles/kg (Fendler et al.). Using carrier-added lutetium-177, the molar activity of our preparations was 0.01 GBq/nmole (0.3 mCi/nmole). Weadministered about 1.45 μCi/5 pmoles, translating to 0.2 nmoles/kg. The masses of PSMA-11 added to the [^177^Lu]-PSMA-617 doses were about 0, 0.2, 4, 20, 40 and 80 nmoles/kg. Based on the molecular weight of PSMA-11 (947 Da), this translates to 0, 0.189, 3.788, 18.94, 37.88 and 75.76 μg/kg, respectively. For a 70 kg man, the administered masses of PSMA-11, in addition to the masses from the [^177^Lu]-PSMA-617 preparation, will be 0, 13.3, 265, 1326, 2652 and 5303 μg, respectively, which is easily achievable. Using these lowered specific activity preparations, we can substantially reduce the uptake in salivary glands and kidneys while maintaining uptake in tumors. The reductions are significant, and if found also in humans they would reduce the severity of side effects such as xerostomia or reduced renal function without compromising tumor uptake. We do realize that data on uptake activity at later time points are needed to concretely establish that the addition of PSMA-11 to the [^177^Lu]-PSMA-617 formulation will reduce non-target organ dose while maintaining the total radiation dose delivered to the tumor. This is beyond the scope of the current manuscript and will be published in a follow-up study. With the reduced uptake in non-target organs, we can potentially treat patients at higher doses or multiple times without causing any serious side effects.

Precedents for such behavior have been observed with small-molecule as well as antibody-based radiopharmaceuticals in pre-clinical or clinical studies. In the pre-clinical studies with Somatostatin-receptor-targeting radiopharmaceuticals, Nicolas et. al. observed that reducing the molar activity by adding a non-radiolabeled ligand from 10 to 2000 pmoles (similar to our study) resulted in significantly reduced uptake of the radiopharmaceutical in the pancreas, salivary gland and marrow without reducing uptake in the tumor xenografts in mice models ^39^. In the clinical studies conducted with PSMA-targeting radiolabeled antibodies (J591) ^9^ or radiolabeled mini-bodies (IA2BM) ^14^, excess cold antibody was administered to minimize liver uptake and under these conditions tumor uptake was maintained even at low molar activities with minimal uptake in salivary glands or kidneys. The radiolabeled antibodies/minibodies were administered along with 10–50 mgs of cold unlabeled molecules, contributing up to 345 nmoles of mAbs (50 mg, average molecular weight 145,000 Da) or 650 nmoles of minibody (50 mg, molecular weight 80,000 Da) to reduce uptake in liver. Kidney uptake with antibodies and minibodies is low because the nephrons in the kidneys restrict the filtration of molecules greater than 60,000 Da in size ^40^. Therefore, we are confident that we can successfully translate our findings to the clinic and improve the pharmacokinetic profile of PSMA TRTs and, ultimately, patient outcomes. With side effects in salivary glands and kidneys minimized, PSMA TRTs could completely alter the treatment landscape of PC, with possible adoption of TRT at earlier stages — even before hormonal and chemotherapy options have been exhausted.

## Conclusions

We demonstrate that by reducing the effective molar activity of our [^177^Lu]-PSMA-617 preparation by addition of cold ligand PSMA-11, we can minimize uptake of the radiopharmaceutical in the salivary glands and kidneys without affecting uptake in the tumor. Because we have reduced the effective molar activity by adding the closely related but non-identical ligand PSMA-11, we believe this method can be easily adapted to several other small-molecule-based PSMA-targeted radiotherapies. Clinical studies are urgently needed to optimize the amount of cold ligand added and to confirm these observations in patients. This simple method has tremendous potential to benefit patients by significantly reducing the dose-limiting toxic side effects of PSMA-targeted therapies and improving outcomes while maintaining quality of life.

## Acknowledgements

Funding support from NIH/NCI R01CA207645-0 (JO, NP), DoD PCRP Idea Award W81XWH-19-1-0536 (NP), NIH/NCI R35 CA232130 (JSL) grants is gratefully acknowledged. Technical and facility services provided by Center of Comparative Medicine & Pathology were supported in part by NIH Grant P30 CA008748.

## Notes

### Competing Interest Statement

The authors have declared no competing interest.

